# Altered thalamo-prefrontal synchrony dynamics during spatial working memory task performance in a SETD1A loss-of-function mouse model of schizophrenia predisposition

**DOI:** 10.64898/2026.01.29.702577

**Authors:** Sofiya Hupalo, David A Kupferschmidt, Ako Ikegami, Mia Railing, Maxym V Myroshnychenko, Gabriel Loewinger, Francisco Pereira, Joseph A Gogos, Joshua A Gordon

## Abstract

Schizophrenia is associated with profound working memory deficits, for which there are no approved treatments. Rare heterozygous null mutations in *SETD1A*, a gene encoding a key epigenetic regulatory protein, have been definitively linked to increased risk for schizophrenia and neurodevelopmental disorders. To investigate how *SETD1A* haploinsufficiency impacts the function of circuits supporting working memory, this study examined neural oscillatory synchrony across a network of brain regions critical for spatial working memory (SWM) in mice carrying a loss-of-function allele in the orthologous *SETD1A* gene. Local field potential recordings were performed in the prefrontal cortex, dorsal and ventral hippocampus, and thalamic nucleus reuniens in male and female wildtype and *Setd1a*^*+/–*^ mice performing a delayed non-match to sample task of SWM. *Setd1a*^*+/–*^ mice exhibited unaltered prefrontal-hippocampal neural oscillatory synchrony across frequencies and task epochs. In contrast, *Setd1a*^*+/–*^ mice displayed reduced beta-frequency synchrony between the prefrontal cortex and nucleus reuniens during SWM maintenance and blunted bidirectional modulation of prefrontal-reuniens beta- and gamma-frequency synchrony across SWM task epochs. Collectively, this work expands our understanding of how genetic risk for schizophrenia alters functional connectivity within distributed circuits supporting SWM.

**GRAPHICAL ABSTRACT:** 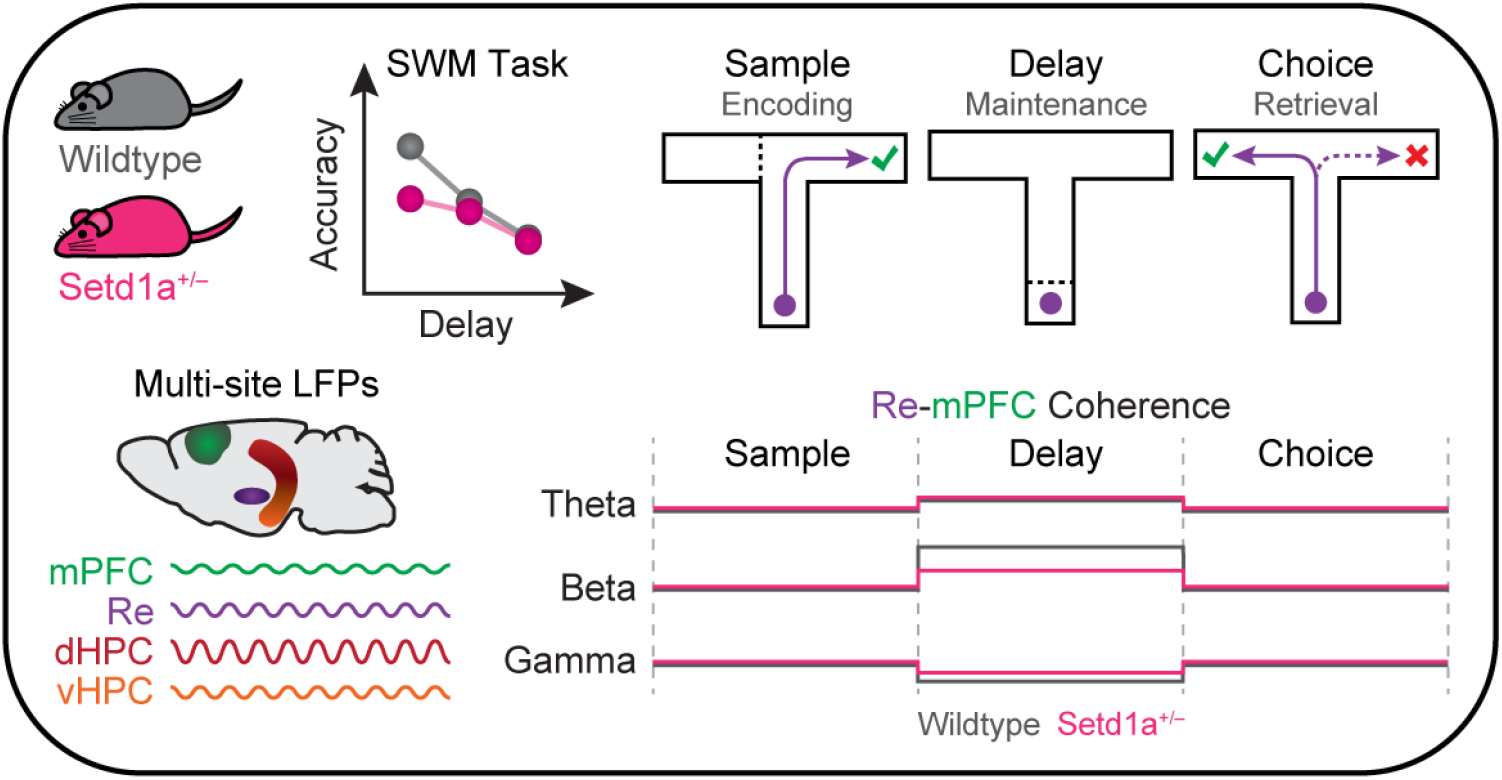

## INTRODUCTION

Working memory—the ability to temporarily maintain and manipulate information to guide behavior—is a core cognitive function. Impairments in this domain are a hallmark of many psychiatric disorders, including schizophrenia [1,2]. Despite working memory deficits being a key predictor of quality of life for individuals with schizophrenia [3–5], no FDA-approved pro-cognitive treatments are currently available [6]. Thus, further research is needed to characterize the neurobiological mechanisms underlying working memory dysfunction in this disorder.

Rare loss-of-function variants in the *SETD1A* gene have emerged as high confidence risk factors for schizophrenia and other neurodevelopmental disorders [7–10]. *SETD1A* encodes a subunit of the histone H3 lysine 4 (H3K4) methyltransferase complex that interacts with active promoters and enhancers to coordinate epigenetic regulation of gene transcription [11]. Transgenic animal models carrying loss-of-function alleles in *SETD1A* provide a powerful framework for linking disease-relevant transcriptional dysregulation to cellular and neural circuit dysfunction. Prior work has demonstrated that *Setd1a* haploinsufficiency alters epigenetic regulation, gene expression, brain development, and the structure and function of the medial prefrontal cortex (mPFC), a region critical for higher-order cognitive processes [12– 16]. Although *Setd1a*^*+/–*^ mice exhibit largely normal gross brain morphology, they show deficits in synaptic transmission, cortical axonal arborization, neuronal density, and dendritic spine density, in regions including the mPFC. Behaviorally, these mice exhibit subtle learning impairments and deficits in working memory and other cognitive domains, with notable phenotypic variation across studies [12,13,15]. Despite extensive evidence for disrupted prefrontal network function in schizophrenia and relevant rodent models [17–19], how *SETD1A* haploinsufficiency alters the coordination of distributed, cognition-modulatory circuits remains unexplored.

Through its interactions with hippocampal and thalamic regions, the rodent mPFC plays a central role in spatial working memory (SWM) [20,21]. Synchronization of neural oscillations in the mPFC, hippocampus, and thalamus facilitates the recruitment of neuronal networks and computations required for successful SWM [22–24]. For example, during *retrieval* of SWM information, theta-frequency (4-12 Hz) synchrony between the mPFC and dorsal hippocampus (dHPC) is elevated [25,26], whereas SWM *encoding* is associated with enhanced coupling between mPFC neuronal firing and ventral hippocampal (vHPC) gamma-frequency (30-70 Hz) oscillations [27]. Inhibition of vHPC terminals in the mPFC during encoding suppresses this gamma synchrony, reduces mPFC encoding of SWM task-relevant information, and impairs SWM performance [27–31]. The *maintenance* of SWM content depends on interactions between the mPFC and the mediodorsal thalamus (MD) [32–36]. Inhibition of MD terminals in the mPFC during the maintenance epoch impairs task performance and reduces beta-frequency (13-30 Hz) MD-mPFC synchrony, whereas similar manipulations during encoding or retrieval have no effect [37]. Lastly, the thalamic nucleus reuniens (Re) supports SWM through its reciprocal connections with the mPFC and HPC [28,38–47]. Beta-frequency synchrony within mPFC-Re-vHPC circuitry is elevated during SWM task performance [48], and Re activity drives mPFC-HPC beta-frequency synchrony while suppressing theta-frequency coupling during memory processing [49], underscoring the capacity of Re to coordinate mPFC-HPC communication. Thus, distinct SWM subprocesses are supported by discrete, frequency-specific modes of synchrony across prefrontal-hippocampal-thalamic circuits.

Here, we characterized neural synchrony within prefrontal-hippocampal-thalamic circuits in *Setd1a*^*+/–*^ mice and their wildtype counterparts during performance of a SWM task. Based on prior studies of other mouse models of genetic risk for schizophrenia [17,18], we hypothesized that *Setd1a*^*+/–*^ mice would exhibit reduced HPC-mPFC synchrony during SWM processing. However, HPC-mPFC synchrony dynamics across the encoding, maintenance, and retrieval epochs were largely unaltered in *Setd1a*^*+/–*^ mice. Instead, *Setd1a*^*+/–*^ mice exhibited blunted bidirectional modulation of Re-mPFC beta- and gamma-frequency synchrony across SWM task epochs, with the most prominent deficit being reduced Re-mPFC beta synchrony during SWM maintenance. Collectively, these findings reveal that *SETD1A* haploinsufficiency gives rise to thalamocortical circuit dysfunction, thereby highlighting that genetic risk factors for schizophrenia yield dissociable patterns of functional dysconnectivity within networks supporting SWM.

## MATERIALS AND METHODS

### Mice

*Setd1a*^tm1a(EUCOMM)Wtsi^ mice (referred to as *Setd1a*^+/–^) were backcrossed for over 10 generations with C57BL/6J mice (The Jackson Laboratory, Bar Harbor, ME) in the laboratory of J. A. Gogos prior to shipment to J. A. Gordon’s laboratory at the NIH. They were maintained by *Setd1a*^*+/–*^ x C57BL/6J crossings. Resulting female and male *Setd1a*^+/–^ and wildtype (WT) littermates (total N = 73; *Setd1a*^+/–^ 16/22, WT 15/20 males/females) were used in the present experiments. Mice were housed in a temperature- and humidity-controlled NIH animal facility on a 12-h light-dark cycle. Except when food-restricted for behavioral training and testing, all mice were given *ad libitum* access to food and water. Pre-surgical mice were group-housed with littermates (up to 5 mice/cage); mice with chronic recording implants were single-housed. All procedures were performed in accordance with the U.S. National Research Council Guide for the Care and Use of Laboratory Animals and were approved by the National Institute of Neurological Disorders and Stroke Animal Care and Use Committee.

### Surgeries

Adult mice were anesthetized with 3-5% isoflurane, placed in a stereotaxic apparatus, and maintained at 0.8-2% isoflurane. Reference and ground wires were adhered to screws placed above the right olfactory bulb (AP: +4.3, M/L: +1.0 mm from bregma) and cerebellum (AP: -6.3), respectively. A tungsten local field potential (LFP) electrode was implanted into the left mPFC (AP: +1.85, ML: -0.4, DV: -1.5 mm below brain surface), dHPC (AP: -2.0, ML: -1.25, DV: -1.35), vHPC (AP: -3.2, ML: -3.5, DV: -3.3), and Re (AP: -0.75, ML: -0.1, DV: -4.0). All wires were connected to a custom electrode interface board fixed to a custom 3D-printed microdrive secured to the skull using dental cement. Prior to the end of surgery, mice were injected with Ketoprofen (10 mg/kg) and Lactated Ringer’s solution (1 mL) to minimize postoperative pain and restore fluid levels. Additional details can be found in the **Supplemental Materials and Methods** section.

### Spatial working memory assay

After recovery from surgery, mice were food-restricted to maintain 85%-90% of their pre-surgery weight. After three days of food restriction and gentle handling, mice underwent two daily sessions of habituation to the T-maze during which they freely visited all arms (baited with milk solution) for 10 min (**Fig. 1a**). Then, mice underwent two consecutive daily shaping sessions (30 trials per session) during which they visited a single available arm (randomized left/right) for milk reward, returned to the start box for reward, and after a 10-sec period visited the alternate single available arm (forced choice) for reward. Mice then began training in the delayed non-match to sample (DNMS) T-maze task. During the sample epoch, mice visited a single available arm for milk reward and returned to the start box. Mice were contained in the start box for a 10-sec delay, after which both goal arms became available, and they had to choose to visit the goal arm not visited during the sample epoch to obtain a reward. Mice performed 10 trials per training session until reaching a criterion of three consecutive days of ≥70% accuracy. Subsequently, mice underwent three daily sessions of “Interleaved testing”, performing 30 trials with delay lengths of 10, 30, or 60 s interleaved randomly within a single session. Mice then performed three daily sessions of “Blocked testing” consisting of 30 trials with consistent delay lengths of 10, 30, or 60 s across sessions; order of the Blocked sessions was counterbalanced across mice. Trials were separated by an intertrial interval of 20 s.

**Figure 1.**
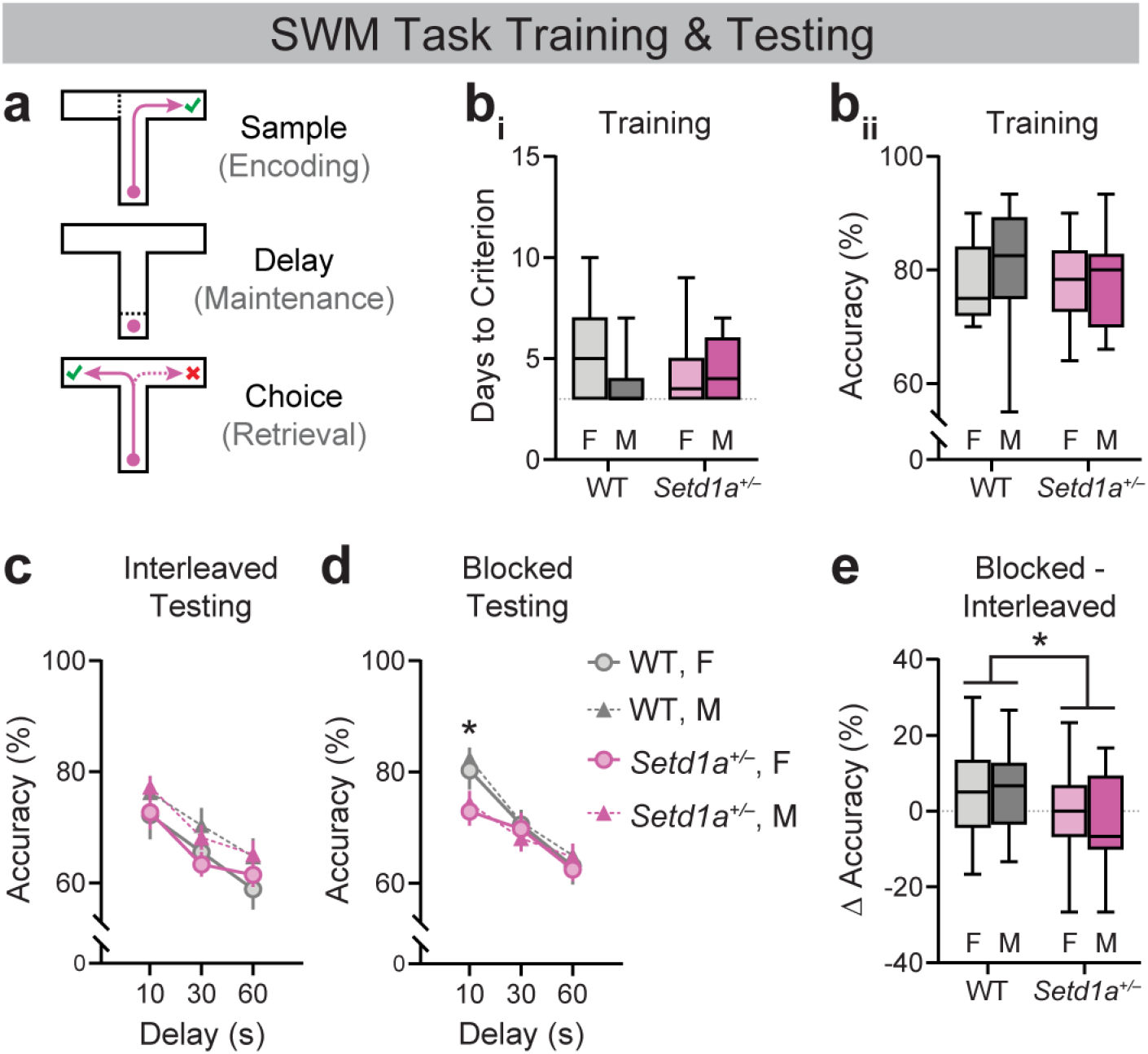
*Setd1a*^*+/–*^ mice exhibit altered SWM task performance with no effect on task acquisition. (**a**) Schematic of trial epochs comprising the delayed non-match-to-sample (DNMS) SWM task. (**b**_**i**_) Days to reach criterion performance of ≥70% correct trials for three consecutive days on the DNMS SWM task in female and male WT and *Setd1a*^*+/–*^ mice (n=15-22); Two-way ANOVA: Main effect of Genotype: *F*(1,69)=0.00, *p*=0.999; Main effect of Sex: *F*(1,69)=2.417, *p*=0.125; Genotype X Sex interaction: *F*(1,69)=1.889, *p*=0.174). (**b**_**ii**_) Average accuracy during training in female and male WT and *Setd1a*^*+/–*^ mice. Two-way ANOVA: Main effect of Genotype: *F*(1,69)=0.569, *p*=0.453; Main effect of Sex: *F*(1,69)=1.209, *p*=0.275; Genotype X Sex interaction: *F*(1,69)=0.727, *p*=0.397). (**c-d**) Accuracy during testing with varying interleaved (within-session) delay lengths (**c**) and blocked (between-session) delay lengths (**d**) in female and male WT and *Setd1a*^*+/–*^ mice. Interleaved: 3-way ANOVA, Main effect of Delay: F(2,136)=36.36, p<0.0001; n=15-22. Blocked: Mixed-effects model, Main effect of Delay: *F*(2,117)=40.20, p<0.0001; Delay x Genotype interaction: *F*(2,117)=3.110, p<0.05; WT vs. *Setd1a*^*+/–*^ at 10-sec delay: *p<0.01, Šídák’s test; n=13-21. (**e**) Difference in performance during 10-sec delay trials between Blocked and Interleaved testing. 2-way ANOVA, Main effect of Genotype: F(1,59)=7.019, *p<0.05).

### Neural data and behavior acquisition

A Cerebus Blackrock Neurotech system acquired electrical recordings through Blackrock Cereplex headstages (μ-series). LFPs referenced to a screw over the olfactory bulb were recorded at 30 kHz, downsampled to 1,500 Hz, and low-pass filtered (≤750 Hz). Video tracking of behaving mice was performed using an Optitrack IR camera (Blackrock Neurotech), ANY-maze software, and an ANY-maze interface providing voltage measures corresponding to the animal’s XY coordinates to analog inputs of the Cerebus system. XY voltages were similarly recorded at 30 kHz and downsampled to 1,500 Hz. Voltage signals were integrated with maze event data in a DataJoint (www.datajoint.com/) database to isolate discrete trials and trial epochs. Of the total 73 mice that completed behavioral testing and LFP recordings, three mice were excluded from subsequent analysis due to improper grounding.

### Coherence estimation

Coherence was estimated on LFP signals downsampled to 1.5 kHz corresponding to each sample run, delay, and choice run epoch and chunked into 1-sec bins. Coherence was estimated from dHPC-mPFC, vHPC-mPFC, vHPC-Re, and Re-mPFC LFP pairs for frequencies ranging from 0-100 Hz using a multi-taper method executed by the Spectral Connectivity function in Python [50]. The time half-bandwidth product was set to 4. Resulting coherence estimates corresponding to the epoch of interest were collapsed across time and subsequently averaged to generate a coherence spectrum per session for every trial epoch.

### Statistical analysis

Statistical analysis was performed using GraphPad Prism and custom Python/R scripts. For behavioral experiments, 2-or 3-way repeated measures ANOVAs were used to assess between-subjects (e.g., genotype, sex) and within-subjects factors (e.g., delay length). In rare instances where a small fraction of datapoints were missing, mixed effects analyses were applied using Prism. Significance (p<0.05) of post hoc comparisons was determined after applying Šídák’s multiple comparisons test. Unless stated otherwise, error bars represent the standard error of the mean and shaded error bands represent 95% confidence intervals.

Functional linear mixed-effects modelling (FLMM) was performed using the fastFMM toolkit [51,52] implemented in R. Individual epoch-level coherence spanning 1-50 Hz was analyzed using three FLMMs. The first modelled coherence as a function of the fixed effect of Genotype with a random intercept for each mouse (coherence ∼ genotype + (1 | mouse_id)). Specifically, for mouse *i*, on epoch *e*, we modelled the coherence at frequency *f* as:

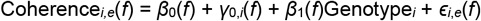

where *ϵ*_*i,e*_(*f*) ∼ N(0, *σ*^2^(*f*)) and *γ*_0,*i*_(*f*) ∼ N(0, *σ*^2^_*γ*_(*f*)). We denoted Genotype_*i*_ = 1 for *Setd1a*^*+/–*^ mice and Genotype_*i*_ = 0 for WT mice. This model was fit to datasets of coherence during sample, delay, or choice epochs of 10-sec delay trials of Blocked testing (**Fig. S2**).

The second modelled coherence as a function of two fixed effects (Genotype and Epoch) and their interaction with a random slope and intercept for Epoch within each mouse (coherence ∼ genotype * epoch + (epoch | mouse_id)):

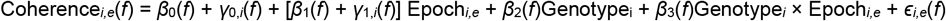

where *ϵ*_*i,e*_(*f*) ∼ N(0, *σ*^2^(*f*)) and *γ*_0,*i*_(*f*) ∼ N(0,Σ_*γ*_(*f*)). We denoted Genotype_*i*_ = 1 for *Setd1a*^*+/–*^ mice and Genotype_*i*_ = 0 for WT mice. We denoted Epoch_*i,e*_ = 0 for sample epochs and Epoch_*i,e*_ = 1 for delay epochs. This model was fit to datasets of coherence during sample and delay epochs of 10-sec delay trials of Blocked testing (**Fig. 2-5**).

**Figure 2.**
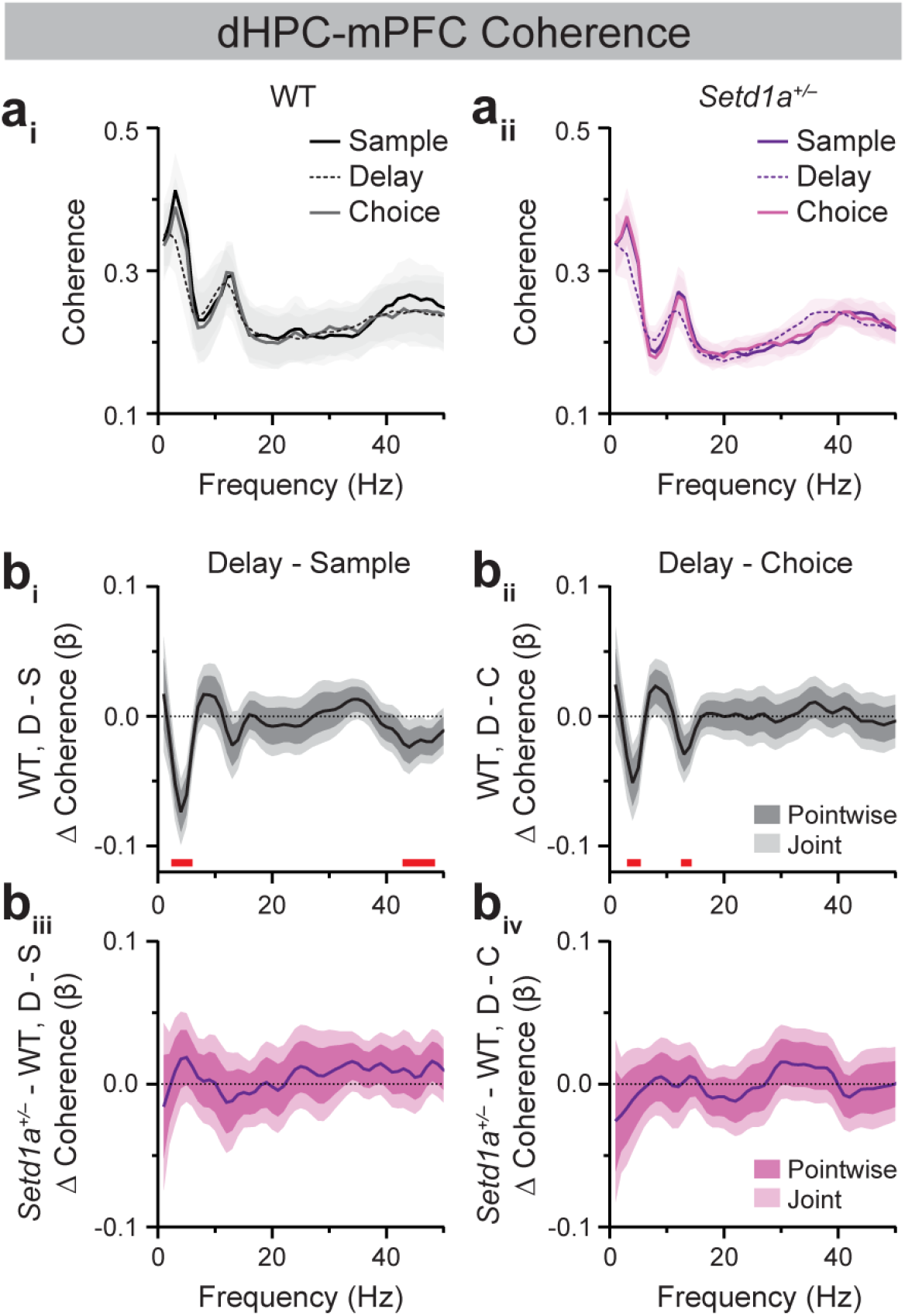
*Setd1a*^*+/–*^ mice exhibit unaltered SWM task epoch-modulated dHPC-mPFC synchrony dynamics. (**a**) Average dHPC-mPFC coherence spectra from WT (**a**_**i**_) and *Setd1a*^*+/–*^ (**a**_**ii**_) mice during sample, delay, and choice epochs of 10-sec delay trials in Blocked testing in the SWM task. (**b**) FLMM of Genotype and Epoch covariate effects on epoch-level dHPC-mPFC coherence data during delay (D) relative to sample (S; **b**_**i**_, **b**_**iii**_) or choice (C; **b**_**ii**_, **b**_**iv**_) epochs. (**b**_**i**_, **b**_**ii**_) Functional coefficient estimates of the Epoch covariate in WT mice, showing differences in the average dHPC-mPFC coherence at each frequency between epoch types. (**b**_**iii**_, **b**_**iv**_) Functional coefficient estimates of the interaction between the Genotype x Epoch covariates, showing how differences in the average dHPC-mPFC coherence at each frequency between epoch types differ between WT and *Setd1a*^*+/–*^ mice. Significant reductions (joint CIs do not contain zero) at specific frequencies are denoted by red bars. n=19 WT, 22 *Setd1a*^*+/–*^ mice.

**Figure 3.**
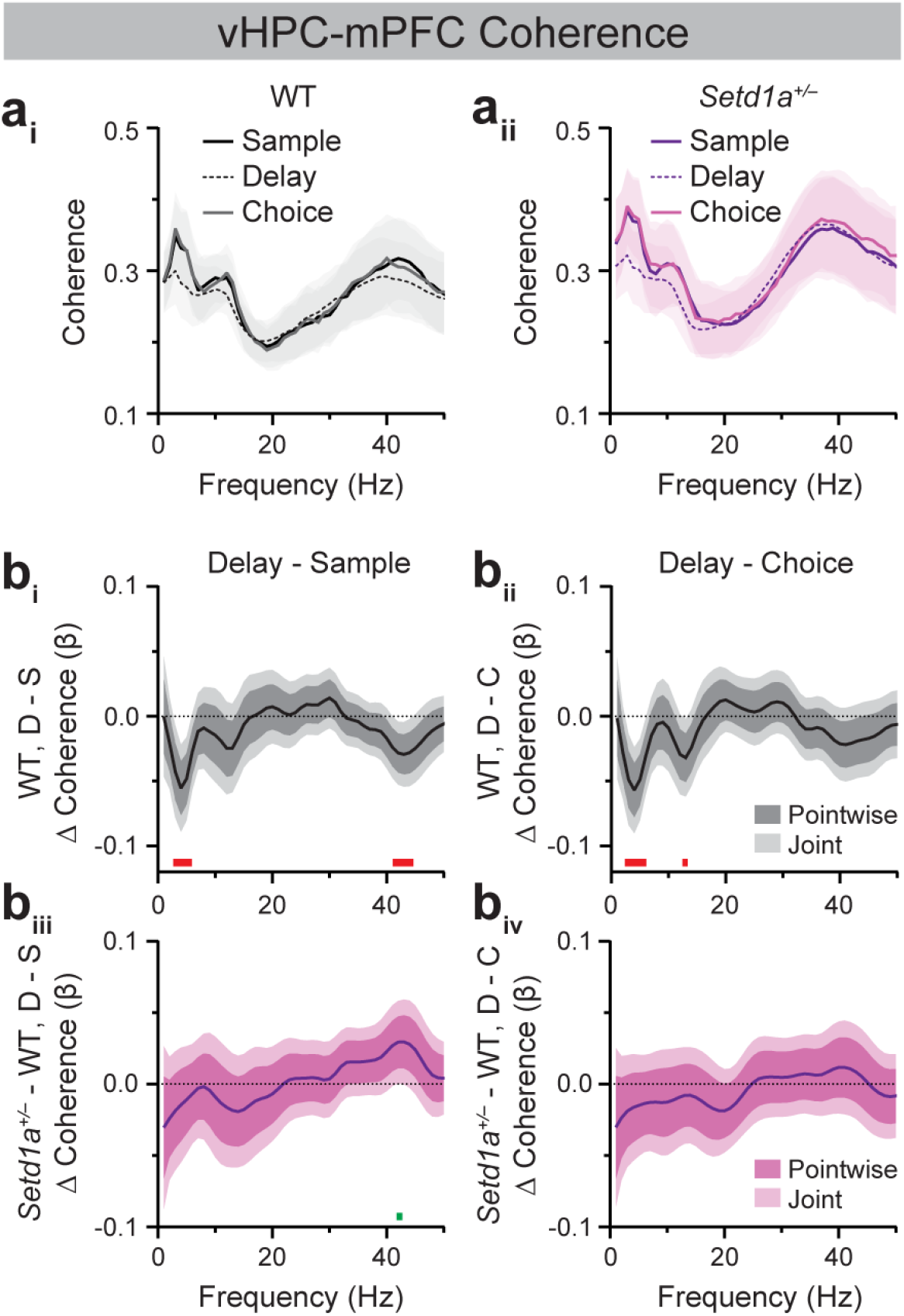
*Setd1a*^*+/–*^ mice exhibit modest blunting of epoch-modulated vHPC-mPFC gamma synchrony dynamics during SWM task performance. (**a**) Average vHPC-mPFC coherence spectra from WT (**a**_**i**_) and *Setd1a*^*+/–*^ (**a**_**ii**_) mice during sample, delay, and choice epochs of 10-sec delay trials in Blocked testing in the SWM task. (**b**) FLMM of Genotype and Epoch covariate effects on epoch-level vHPC-mPFC coherence data during delay (D) relative to sample (S; **b**_**i**_, **b**_**iii**_) or choice (C; **b**_**ii**_, **b**_**iv**_) epochs (**b**_**i**_, **b**_**ii**_) Functional coefficient estimates of the Epoch covariate in WT mice, showing differences in the average vHPC-mPFC coherence at each frequency between epoch types. (**b**_**iii**_, **b**_**iv**_) Functional coefficient estimates of the interaction between the Genotype x Epoch covariates, showing how differences in the average vHPC-mPFC coherence at each frequency between epoch types differ between WT and *Setd1a*^*+/–*^ mice. Significant enhancements and reductions (joint CIs do not contain zero) at specific frequencies are denoted by green and red bars, respectively. n=16 WT, 20 *Setd1a*^*+/–*^ mice.

**Figure 4.**
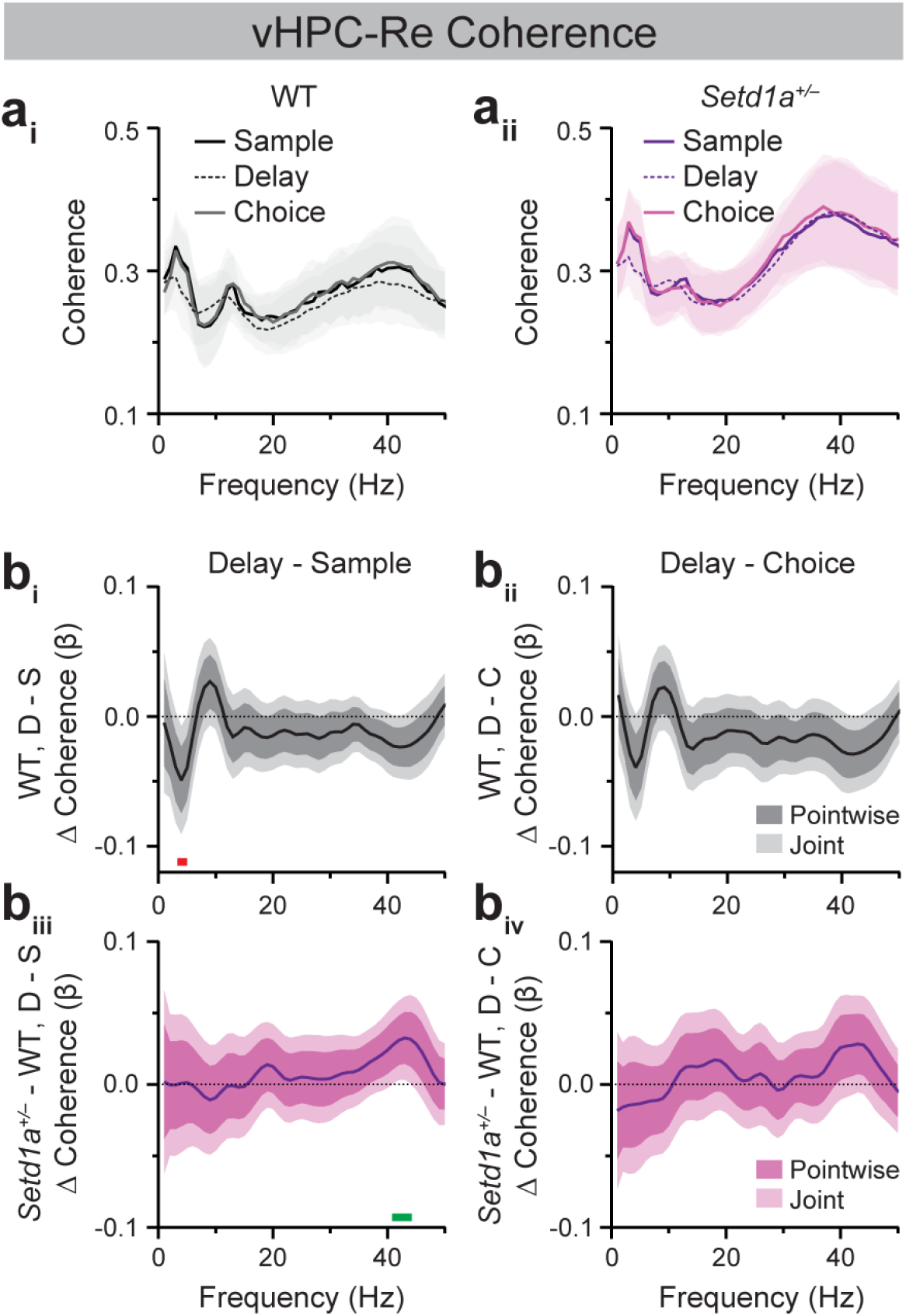
*Setd1a*^*+/–*^ mice exhibit modest blunting of epoch-modulated vHPC-Re gamma synchrony dynamics during SWM task performance. (**a**) Average vHPC-Re coherence spectra from WT (**a**_**i**_) and *Setd1a*^*+/–*^ (**a**_**ii**_) mice during sample, delay, and choice epochs of 10-sec delay trials in Blocked testing in the SWM task. (**b**) FLMM of Genotype and Epoch covariate effects on epoch-level vHPC-Re coherence data during delay (D) relative to sample (S; **b**_**i**_, **b**_**iii**_) or choice (C; **b**_**ii**_, **b**_**iv**_) epochs. (**b**_**i**_, **b**_**ii**_) Functional coefficient estimates of the Epoch covariate in WT mice, showing differences in the average vHPC-Re coherence at each frequency between epoch types. (**b**_**iii**_, **b**_**iv**_) Functional coefficient estimates of the interaction between the Genotype x Epoch covariates, showing how differences in the average vHPC-Re coherence at each frequency between epoch types differ between WT and *Setd1a*^*+/–*^ mice. Significant enhancements and reductions (joint CIs do not contain zero) at specific frequencies are denoted by green and red bars, respectively. n=14 WT, 25 *Setd1a*^*+/–*^ mice.

**Figure 5.**
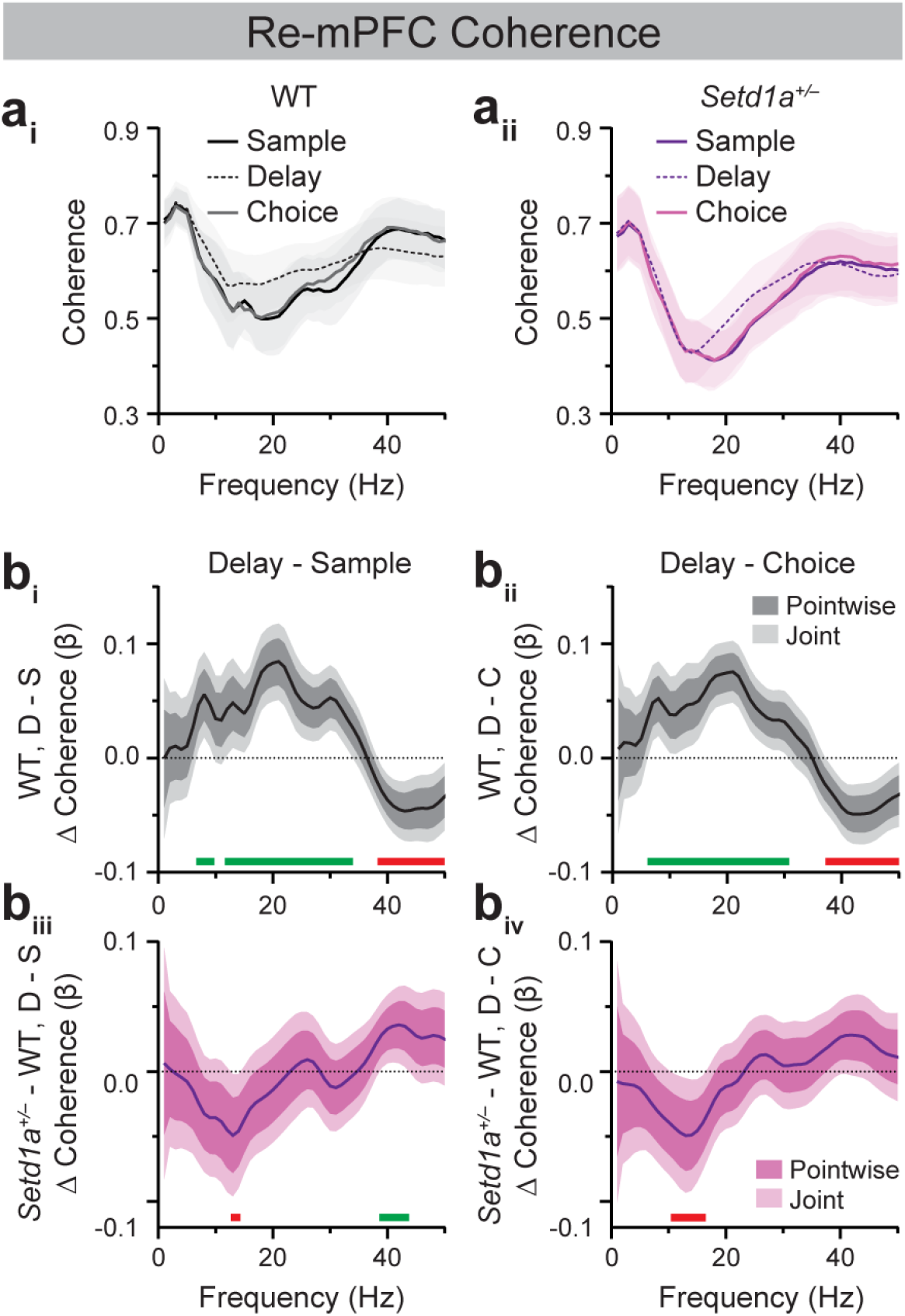
Bidirectional modulation of Re-mPFC beta and gamma synchrony across SWM task epochs is blunted in *Setd1a*^*+/–*^ mice. (**a**) Average Re-mPFC coherence spectra from WT (**a**_**i**_) and *Setd1a*^*+/–*^ (**a**_**ii**_) mice during sample, delay, and choice epochs of 10-sec delay trials in Blocked testing in the SWM task. (**b**) FLMM of Genotype and Epoch covariate effects on epoch-level Re-mPFC coherence data during delay (D) relative to sample (S; **b**_**i**_, **b**_**iii**_) or choice (C; **b**_**ii**_, **b**_**iv**_) epochs. (**b**_**i**_, **b**_**ii**_) Functional coefficient estimates of the Epoch covariate in WT mice, showing differences in the average Re-mPFC coherence at each frequency between epoch types. (**b**_**iii**_, **b**_**iv**_) Functional coefficient estimates of the interaction between the Genotype x Epoch covariates, showing how differences in the average Re-mPFC coherence at each frequency between epoch types differ between WT and *Setd1a*^*+/–*^ mice. Significant enhancements and reductions (joint CIs do not contain zero) at specific frequencies are denoted by green and red bars, respectively. n=19 WT, 23 *Setd1a*^*+/–*^ mice.

The third model was identical to the second, except that we denoted Epoch_*i,e*_ = 0 for choice epochs and Epoch_*i,e*_ = 1 for delay epochs. This model was fit to datasets of coherence during choice and delay epochs of 10-sec delay trials of Blocked testing (**Fig. 2-5**).

Prior to selection, models were tested using different random effects structures; models that successfully converged and yielded lower Akaike and Bayesian Information Criteria (AIC and BIC) values were selected. FLMM generates Pointwise and Joint 95% confidence intervals (CIs) around the resulting functional coefficients. Frequencies at which Pointwise 95% CIs do not contain Y = 0 indicate individual frequencies of statistical significance, without correction for multiple comparisons. Joint 95% CIs, generated by exploiting the correlation between neighboring frequencies, are adjusted for multiple comparisons; thus, frequencies at which Joint 95% CIs do not contain 0 are interpreted as statistically significant.

## RESULTS

### *Setd1a*^+/–^ mice show altered SWM task performance with no impairment in task acquisition

Male and female *Setd1a*^+/–^ mice and WT controls (n=16 male *Setd1a*^+/–^, 22 female *Setd1a*^+/–^, 15 male WT, and 20 female WT) were trained and tested on the DNMS task of SWM. Each trial consisted of three epochs: a “sample” epoch wherein mice must encode spatial information as they navigate to a maze arm, a “delay” epoch over which mice must maintain that information, and a “choice” epoch wherein mice must retrieve that information to navigate to the opposite arm to obtain reward (**Fig. 1a**; e.g., [17,53]). Using a similar task, Mukai and colleagues [12] reported that male *Setd1a*^*+****/–***^ mice acquire the task at rates comparable to controls but exhibit impaired performance when tested in sessions with trials of mixed delay lengths. Extending and largely consistent with these findings, we found no statistically significant differences between *Setd1a*^*+****/–***^ and WT mice in the number of sessions required to reach training criterion (**Fig. 1b_i_**; Main effect of Genotype: *F*(1,69)=0.00, *p*=0.999). Average accuracy across training sessions was also indistinguishable between the two groups (**Fig. 1b_ii_**; Main effect of Genotype: *F*(1,69)=0.569, *p*=0.453). No significant effects of sex or sex-by-genotype interactions were observed.

During Interleaved (**Fig. 1c**) and Blocked testing (**Fig. 1d**), average accuracy declined in trials with longer delay lengths, consistent with the DNMS task being an assay of SWM. As during training, no sex differences were observed during testing. *Setd1a*^+/–^ mice performed comparably to WT controls in Interleaved testing, in which delay lengths varied unpredictably within a session (**Fig. 1c**; Main effect of Delay: *F*(2,136)=36.36, *p*<0.0001). In contrast, *Setd1a*^+/–^ mice exhibited reduced accuracy relative to controls during 10-sec delay trials in Blocked sessions, where the delay was held constant across trials (**Fig. 1d**; Main effect of Delay: *F*(2,117)=40.20, *p*<0.0001; Delay x Genotype interaction: *F*(2,117)=3.110, *p*<0.05; WT vs. Setd1a^+/–^ at 10-sec delay: \**p*<0.01, Šídák’s test). This genotype-dependent difference was driven by greater performance improvement in WT mice during 10-sec trials of Blocked relative to Interleaved testing, consistent with enhanced performance under lower cognitive demand (**Fig. 1e**; Main effect of Genotype: *F*(1,59)=7.019, \**p*<0.05). Together, these findings suggest that *Setd1a*^**+/–**^ mice exhibit modest SWM task performance deficits that emerge under conditions of reduced cognitive demand.

### WT and *Setd1a*^+/–^ mice display comparable hippocampal-prefrontal coherence during SWM task performance

To assess SWM task-relevant neural synchrony dynamics and their potential alteration by *SETD1A* haploinsufficiency, we analyzed inter-regional coherence using LFP recordings from the mPFC, dHPC, vHPC, and Re in WT and *Setd1a*^+/–^ mice performing the DNMS task (**Fig. S1**). Our analyses focused on 10-sec trials from Blocked testing sessions, the condition under which *Setd1a*^+/–^ mice displayed performance deficits. We first quantified broadband oscillatory synchrony using mouse-averaged LFP-LFP coherence between dHPC-mPFC and vHPC-mPFC across DNMS task epochs (**Fig. 1a**).

These analyses revealed comparable hippocampal-prefrontal synchrony dynamics in WT and *Setd1a*^+/–^ mice (**Fig. 2a, 3a**). To quantify potential differences in coherence while accounting for trial-to-trial variability and inter-animal heterogeneity, we used FLMM, which enables hypothesis testing at every frequency in a frequency-unbiased manner [52]. FLMM of epoch-specific coherence, with significance testing using Joint 95% Cis, showed no genotype-dependent differences in dHPC-mPFC (**Fig. S2a_ii, iv, vi_**) or vHPC-mPFC (**Fig. S2b_ii, iv, vi_**) coherence during the sample, delay, or choice epochs at any frequency between 1-50 Hz.

Because hippocampal-prefrontal synchrony is modulated by the different cognitive subprocesses and behavioral states comprising a SWM task trial, we assessed genotype-dependent differences across DNMS task epochs using FLMM of coherence as a function of genotype, epoch, and their interaction. In WT mice, dHPC-mPFC (**Fig. 2b_i, ii_**) and vHPC-mPFC (**Fig. 3b_i, ii_**) coherence in the lower theta-frequency range was significantly reduced during the delay epoch relative to the sample and choice epochs, consistent with the established association between HPC-mPFC theta synchrony and active navigation (Delay vs. Sample, dHPC-mPFC: ∼3-6 Hz; vHPC-mPFC: ∼3-5 Hz; Delay vs. Choice, dHPC-mPFC: ∼4-5 Hz, ∼13-14 Hz; vHPC-mPFC: ∼3-6 Hz, ∼13 Hz; [54]). WT mice also exhibited reduced dHPC-mPFC (**Fig. 2b_i_**, ∼43-48 Hz) and vHPC-mPFC gamma synchrony (**Fig. 3b_i_**, ∼42-44 Hz) during the delay relative to the sample epoch, consistent with prior evidence that vHPC-mPFC gamma synchrony supports encoding of spatial information into WM [27]. Analysis of Genotype x Epoch interactions revealed that dHPC-mPFC coherence was modulated similarly across epochs in WT and *Setd1a*^+/–^ mice (**Fig. 2b_iii, iv_**). Similarly, *Setd1a*^+/–^ mice showed indistinguishable modulation of vHPC-mPFC coherence across DNMS task epochs relative to WT mice, except for a subtle attenuation in *Setd1a*^+/–^ mice of the decrease in gamma coherence in the delay relative to sample epoch seen in WT mice at ∼42 Hz (**Fig. 3b_iii_**).

### *Setd1a*^+/–^ mice display modest alterations in vHPC-Re coherence during SWM task performance

vHPC-Re coherence dynamics during DNMS performance resembled those of the dHPC-mPFC and vHPC-mPFC pairs (**Fig. 4a**). In WT mice, vHPC-Re theta (∼4 Hz) coherence was significantly reduced in the delay relative to the sample epoch (**Fig. 4b_i_**). No significant differences were observed between WT and *Setd1a*^+/–^ mice in sample-, delay-, or choice-related vHPC-Re coherence at any frequency between 1-50 Hz (**Fig. S2c**). However, as with vHPC-mPFC coherence, the difference in delay vs. sample vHPC-Re gamma coherence (∼41-44 Hz) was significantly altered in *Setd1a*^+/–^ mice relative to controls (**Fig. 4b_iii_**).

### Re-mPFC beta- and gamma-frequency coherence is bidirectionally modulated across SWM trial epochs

Next, we examined Re-mPFC neural synchrony dynamics during SWM (**Fig. 5a_i_**). Using frequency-unbiased FLMM to assess how Re-mPFC coherence changed across DNMS task epochs in WT mice, we observed that Re-mPFC theta and beta coherence was significantly elevated during the delay epoch relative to sample and choice (**Fig. 5b_i, ii_**). These delay-related increases spanned very comparable frequency ranges for the delay vs. sample (∼7-9 Hz, ∼12-33 Hz) and delay vs. choice (∼7-30 Hz) comparisons. Notably, delay-related increases in beta coherence were unique to the Re-mPFC circuit (**Figs. 2-4b_i, ii_**). In contrast to beta coherence, Re-mPFC gamma coherence was significantly reduced during the delay epoch relative to the sample and choice epochs (**Fig. 5b_i, ii_**; Delay vs. Sample: ∼39-50 Hz; Delay vs. Choice: ∼38-50 Hz). Together, these results demonstrate a robust, bidirectional modulation of Re-mPFC synchrony across SWM task epochs, characterized by enhanced theta/beta inter-regional coupling and suppressed gamma coupling during working memory maintenance.

### *Setd1a*^+/–^ mice exhibit altered Re-mPFC coherence dynamics across DNMS trial epochs

Lastly, we assessed genotype-dependent Re-mPFC synchrony dynamics (**Fig 5a_ii_**). FLMM revealed that *Setd1a*^+/–^ mice displayed reduced delay-related Re-mPFC beta (∼12-18 Hz) coherence relative to controls (**Fig. S2d_iv_**), with no genotype differences during the sample (**Fig. S2d_ii_**) or choice epochs (**Fig. S2d_vi_**). By analyzing Analysis of Genotype x Epoch interactions further showed that compared to WT mice, *Setd1a*^+/–^ mice displayed blunted enhancement of delay-related Re-mPFC beta coherence relative to that seen in the sample and choice epochs (**Fig. 5b_iii, iv_**). These genotype-dependent effects were restricted to narrow beta-frequency ranges (∼13-14 Hz for the delay vs. sample comparison; ∼11-16 Hz for the delay vs. choice). In addition to beta-band alterations, *Setd1a*^+/–^ mice also exhibited blunted Re-mPFC gamma coherence increases during the sample relative to the delay epoch (∼39-43 Hz) compared to WT controls (**Fig. 5b_iii_**).

Together, these findings demonstrate that SETD1A haploinsufficiency selectively disrupts the bidirectional, epoch-specific modulation of Re-mPFC beta- and gamma-frequency synchrony during SWM task performance, with the most prominent deficit being reduced Re-mPFC beta synchrony during SWM maintenance.

## DISCUSSION

The current studies characterized neural synchrony dynamics across prefrontal-hippocampal-thalamic circuits in WT and *SETD1A* haploinsufficient mice modeling genetic risk for schizophrenia. Consistent with prior studies, WT mice exhibited elevated HPC-mPFC theta synchrony during the sample and choice epochs of the SWM task, which coincide with periods of active navigation (**Fig. 2, 3**; [25,54]). vHPC-mPFC gamma synchrony was also enhanced during the sample relative to the delay epoch, corroborating previous findings demonstrating a role for this circuit in SWM encoding (**Fig 3**; [27]).

Extending prior work, these studies characterized Re synchrony with the mPFC and HPC during SWM performance. Similar to vHPC-mPFC dynamics, vHPC-Re theta coherence was elevated during the sample relative to the delay epoch of the DNMS task (**Fig. 4**). Within the Re-mPFC circuit, synchrony was bidirectionally modulated across SWM epochs, with theta- and beta-frequency coupling enhanced during the delay epoch and gamma-frequency coupling enhanced during the sample and choice epochs (**Fig. 5**). Together, these findings provide new insight into the dynamic, frequency-specific coordination of Re-mPFC circuitry during SWM.

*Setd1a*^*+/–*^ mice acquired the DNMS task comparably to WT controls and did not display performance deficits when delay durations varied unpredictably within a session. However, *Setd1a*^*+/–*^ mice performed worse than controls during 10-sec delay trials of Blocked sessions, in which the delay duration was held constant across trials. Together, these findings suggest that *Setd1a*^*+/–*^ mice display relatively subtle SWM impairments that may reflect difficulty adopting a stable performance strategy under consistent delay demands. Although broadly aligned with prior reports, these results contrast with reports of more pronounced SWM and other cognitive deficits in *Setd1a*^*+/–*^ mice [12,13,15]. Such cross-study differences may arise from variability in experimental design, including the specific SWM test used (e.g., Y-maze, delayed alternation, delay non-match to sample/place), mouse age and sex, the loss-of-function mutation used to induce haploinsufficiency, and the extent of training and testing (e.g., number of trials/session). Additionally, these differences may reflect inherent variability in the expression of modest behavioral phenotypes in *Setd1a*^*+/–*^ mice. Notably, similar variability has been reported in *Df(16)A*^+/–^ mice modeling the schizophrenia-predisposing 22q11.2 microdeletion, which exhibit inconsistent magnitudes of DNMS task learning deficits across studies [17][18]. Consistently across studies, these learning deficits were associated with robust decreases in theta and gamma HPC-mPFC synchrony. Together, this evidence suggests that genetic risk for schizophrenia may yield more robust and reliable alterations at the cellular and circuit levels than at the level of behavior.

Given that *Df(16)A*^+/–^ mice display reduced HPC-mPFC theta and gamma synchrony during SWM performance, we hypothesized that *Setd1a*^+/–^ mice would exhibit similar alterations. However, SWM-related dHPC-mPFC and vHPC-mPFC synchrony did not differ between WT and *Setd1a*^+/–^ mice (**Fig. 2, 3**). Instead, *Setd1a*^+/–^ mice displayed modest blunting of SWM epoch-modulated vHPC-Re gamma synchrony increases in the sample relative to the delay epoch (**Fig. 4**).

Moreover, these mice showed reduced delay-related Re-mPFC beta synchrony and a blunted increase in beta coupling during the delay epoch compared to the sample and choice epochs (**Fig. S2d, 5**). In addition, *Setd1a*^+/–^ mice displayed blunted Re-mPFC gamma synchrony during the sample relative to the delay epoch (**Fig. 5**). Combined, these finding demonstrate that bidirectional modulation of Re-mPFC beta and gamma synchrony during distinct SWM task epochs is disrupted in *Setd1a*^+/–^ mice.

It remains unclear whether reduced delay-related Re-mPFC beta coherence in *Setd1a*^+/–^ mice causally contributes to SWM impairment or instead reflects broader physiological processes unrelated to cognition. This uncertainty stems in part from an incomplete understanding of how bidirectional Re-mPFC projections support SWM performance and whether beta synchrony itself plays a causal role. Across species, beta oscillatory activity within thalamo-prefrontal circuits has been implicated in top-down control and motor output during goal-directed behavior. In rodents, beta synchrony between the mPFC and MD thalamus associates with successful maintenance of SWM task-relevant spatial information across a delay [35,37]. Moreover, beta-frequency synchrony within mPFC-Re-vHPC circuits is elevated during SWM task performance relative to a non-SWM task [48]. In non-human primates, transient prefrontal bursts of alpha- and beta-frequency activity are associated with top-down informational processing and are anti-correlated with gamma activity, which is thought to represent reflect the maintenance of sensory information held in WM [24,55]. In the current study, the bidirectional modulation of Re-mPFC theta/beta and gamma synchrony across SWM task epochs in control mice— characterized by greater theta/beta synchrony during delay and enhanced gamma synchrony during sample and choice epochs—is aligned with this literature. Notably, cortical beta bursts have also been linked to suppression of motor activity [56], consistent with our observation that Re-mPFC beta synchrony is elevated during the delay epoch when animals are relatively stationary. Future studies that directly manipulate beta-frequency interactions within prefrontal-hippocampal-thalamic circuits will be essential for determining their causal contributions to both intact and disordered SWM.

## DATA AVAILABILITY

All data are available in the main text or the Supplemental Material.

## ACKNOWLEDGEMENTS

We thank all members of the Gordon laboratory for their generous discussions, feedback, and technical advice. In particular, we thank Dr. Johannes Passecker for behavioral and electrophysiology analysis expertise, and Emily Alway, Kirstin Gilchrist, and Thomas Clarity for assistance with behavioral profiling of the *Setd1a*^*+/–*^ mice. We thank George Dold and colleagues in the NIMH Section on Instrumentation for their expert design and support of the T-maze apparatus. We thank Dr. Vincent Schram and colleagues in the NICHD Imaging Core for their training and assistance in confocal imaging. We thank the Porter Neuroscience Research Center Animal Care Staff for their exceptional animal husbandry.

## AUTHOR CONTRIBUTIONS

Conceptualization, S.H., D.A.K., J.A.G.^2^

Methodology, S.H., D.A.K., M.R., M.V.M., G.L., J.A.G.^1^, J.A.G.^2^

Investigation and data analysis, S.H., D.A.K., A.I., M.R., M.V.M. Funding acquisition, S.H., J.A.G.^2^

Project administration, S.H., D.A.K. Supervision, D.A.K., J.A.G.^2^ Writing (original draft), S.H., D.A.K.

Writing (review & editing), S.H., D.A.K., A.I., M.R., G.L., J.A.G.^1^, J.A.G.^2^

## FUNDING

This work was supported by the Division of Intramural Research of the NINDS (ZIA NS003168) and the NIGMS Postdoctoral Research Associate Training (PRAT) Program Fellowship (FI2-GM133387, SH).

## COMPETING INTERESTS

The authors have nothing to disclose.

## SUPPLEMENTAL MATERIAL

### Surgeries

Adult mice (2.5-4 months) were anesthetized with 3-5% isoflurane (v/v in oxygen, flow rate of 1 L/min) and placed in a stereotaxic apparatus (Kopf Instruments) using non-rupture ear bars. Mice were maintained at 0.8-2% isoflurane on a 45°C heating pad for the duration of the surgery. Prior to initiating and periodically throughout the surgery, anesthesia depth was tested by a toe pinch; mice with any response were given supplemental isoflurane anesthesia. Tips of all surgical instruments were sterilized in a hot bead sterilizer and cooled prior to surgery.

The scalp was opened via a 2-cm excision, skin was retracted, and 0.5-1 mm burr holes were drilled in the skull over the olfactory bulb and cerebellum, as well as the left mPFC, dHPC, vHPC, and Re (Franklin & Paxinos, 2008). Skull screws were imbedded over the right olfactory bulb (AP: +4.3, M/L: +1.0 mm from bregma) and cerebellum (AP: -6.3). A tungsten wire local field potential (LFP) electrode (76-μm diameter; California Fine Wire) coated with DiI lipophilic stain (ThermoFisher) was implanted into the left mPFC (AP: +1.85, ML: -0.4, DV: -1.5 mm below brain surface), dHPC (AP: - 2.0, ML: -1.25, DV: -1.35 from brain surface), vHPC (AP: -3.2, ML: -3.5, DV: -3.3 from brain surface), and Re (AP: -0.75, ML: -0.1, DV: -4.0 from brain surface). Reference and ground wires were adhered to screws placed above the olfactory bulb and cerebellum, respectively. All wires were connected to a custom electrode interface board (printed circuit board, MyroPCB; solder paste, Digi-Key; solder stencil, Pololu; connector, Omnetics; gold pins, Neuralynx) fixed to a custom 3D-printed microdrive secured via dental cement (combination of Metabond/RelyX Unicem/Filtek, Henry Schein) to the skull surface. The microdrive was enclosed in a 3D-printed shield to protect its components. Vetbond tissue adhesive (Fisher Scientific) or Vicryl absorbable sutures (size 5-0, using simple interrupted stitches) were used as necessary to bring the incision into contact with the shield. 30 min prior to the end of surgery, mice were injected with warmed Ketoprofen (10 mg/kg, s.c.) in Lactated Ringer’s solution, and additional Lactated Ringer’s solution (1 mL), to minimize postoperative pain and restore fluid levels.

Mice recovered in a clean homecage with food pellets and/or a rodent diet gel pack. The recovery cage was placed half on/off a heating pad to allow mice to regulate their temperature during the initial recovery period. Ketoprofen (10 mg/kg, s.c.) in Lactated Ringer’s solution was administered 24- and 48-h post-surgery for postoperative pain management. Mice recovered for a minimum of 10 days before undergoing food restriction for behavioral experiments.

### Spatial working memory assay

#### Maze apparatus

A custom-built automated T-maze was used for assessing SWM performance. Each arm of the T-maze was 12.7 cm wide and 30.5 cm high. The center arm (stem) was 60-cm long (including an 18-cm start box), and the goal arms were 26.7-cm long. The start box and goal arms contained a reward port for delivery of sweetened condensed milk (∼40 μl, 20% in deionized water) through a blunt 20G needle. Delivery was triggered when mice broke infrared beams positioned in the maze walls at task-relevant periods. A circular rotary platform, positioned such that one half comprises the choice point of the maze, made 180° rotations during the delay epoch of each trial to prevent mice from using scent/path cues to guide their subsequent choice (described below). Maze components were controlled using custom Python scripts. Maze events (inputs and outputs) were timestamped using Python-mediated text comments sent to Blackrock Central software.

#### DNMS task training and testing

At least 10 days post-surgery, mice were food-restricted diet to maintain 85%–90% of their body weight (after correcting for weight of the implanted microdrive). After three days of food restriction and gentle handling, mice underwent two daily sessions of habituation to the T-maze during which they freely visited all arms (baited with milk solution) for 10 min (**Fig. 1a**). Mice were tethered to the electrical cable on the second habituation session and all sessions thereafter. Mice underwent two consecutive daily shaping sessions during which they visited a single available arm (randomized left/right) for milk reward, returned to the start box for reward, and after a 10-sec period visited the alternate single available arm (forced choice) for reward. Shaping sessions consisted of up to 30 trials over 30 min.

Mice then began training in the delayed non-match to sample (DNMS) task. Each DNMS trial consisted of 3 epochs: Sample, Delay, and Choice. In the sample epoch, mice visited a single available arm (randomized between left or right arms) for milk reward and returned to the start box. Mice were contained in the start box for a 10-sec delay period, after which both goal arms became available, and they had to choose to visit the goal arm not visited during the sample epoch to obtain a reward. Mice performed 10 trials per daily training session until reaching a criterion of three consecutive days of ≥70% correct trials. Subsequently, mice underwent three daily sessions of “Interleaved testing”, performing 30 trials with delay lengths of 10, 30, or 60 s interleaved randomly within a single session. Mice then performed three daily sessions of “Blocked testing” consisting of 30 trials with consistent delay lengths of 10, 30, or 60 s across sessions; order of the Blocked sessions was counterbalanced across mice. Trials were separated by an intertrial interval of 20 s. All behavioral experiments were performed during the light cycle. Experimenters conducting behavior training and testing were blinded to mouse genotype.

### Histology

Mice were deeply anesthetized with sodium pentobarbital (FATAL-PLUS, 20 mg/kg, i.p.) or isoflurane (5% in oxygen) and transcardially perfused with 10 mL PBS followed by 10 mL 4% paraformaldehyde in PBS. Brains were post-fixed in 4% paraformaldehyde at 4°C for 24-48 h and cryoprotected in 30% phosphate-buffered sucrose for 3 days at 4°C. Brains were sectioned (40 μm) using a cryostat and mounted with DAPI Fluoromount-G mounting medium (Southern Biotech). Electrode placements were verified by examining DiI fluorescence from the stain coating each LFP wire. Only data from verified recording sites were used for coherence analyses.

**Figure S1.**
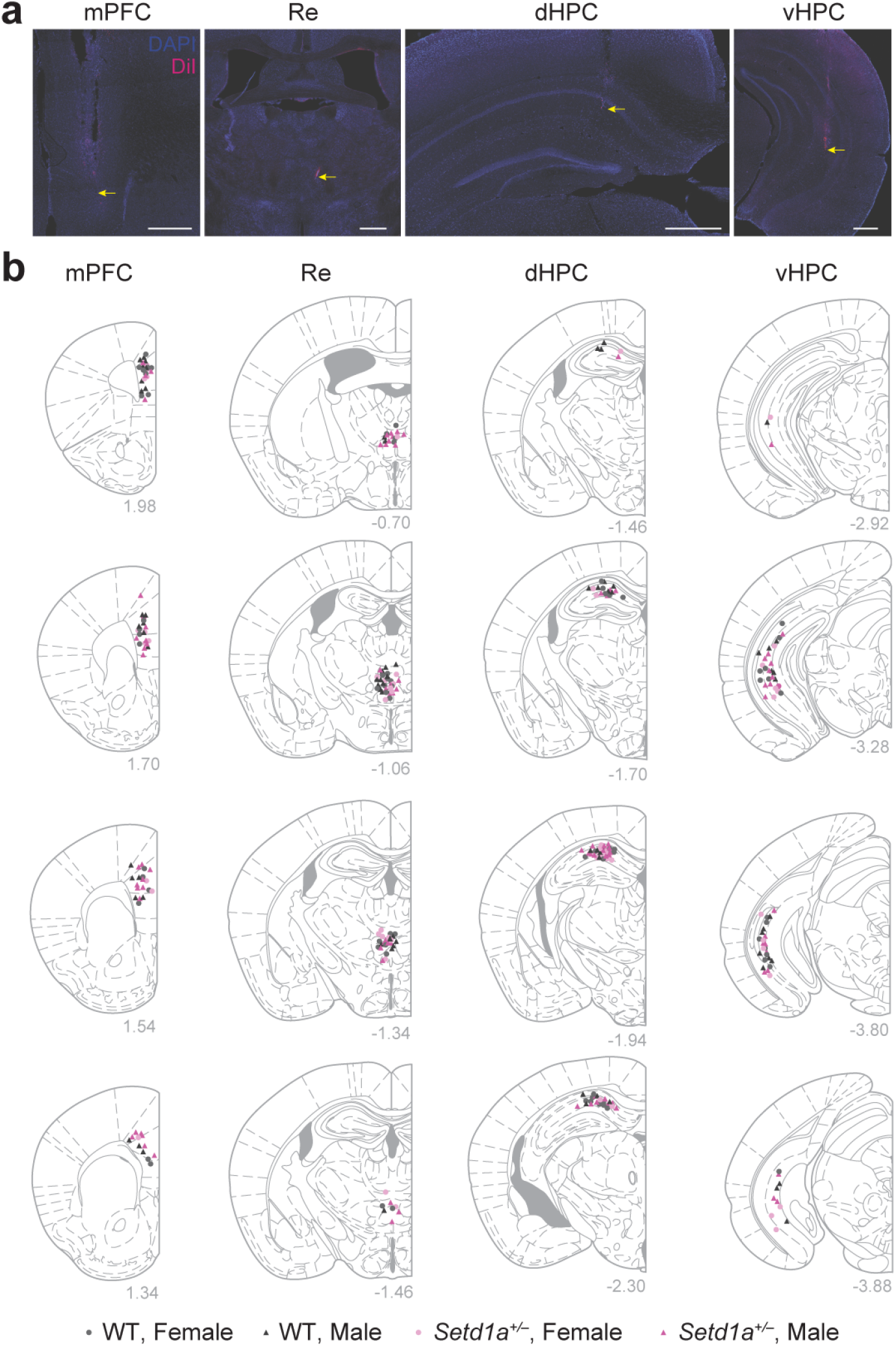
Histology of electrode implant sites. (**a**) Representative histology showing electrode placements in mPFC, vHPC, dHPC, and Re. Yellow arrows indicate the estimated electrode termination sites. Scale bars = 500μm. (**b**) Electrode placements in mPFC, vHPC, dHPC, and Re of female and male WT and *Setd1a*^*+/–*^ mice.

**Figure S2.**
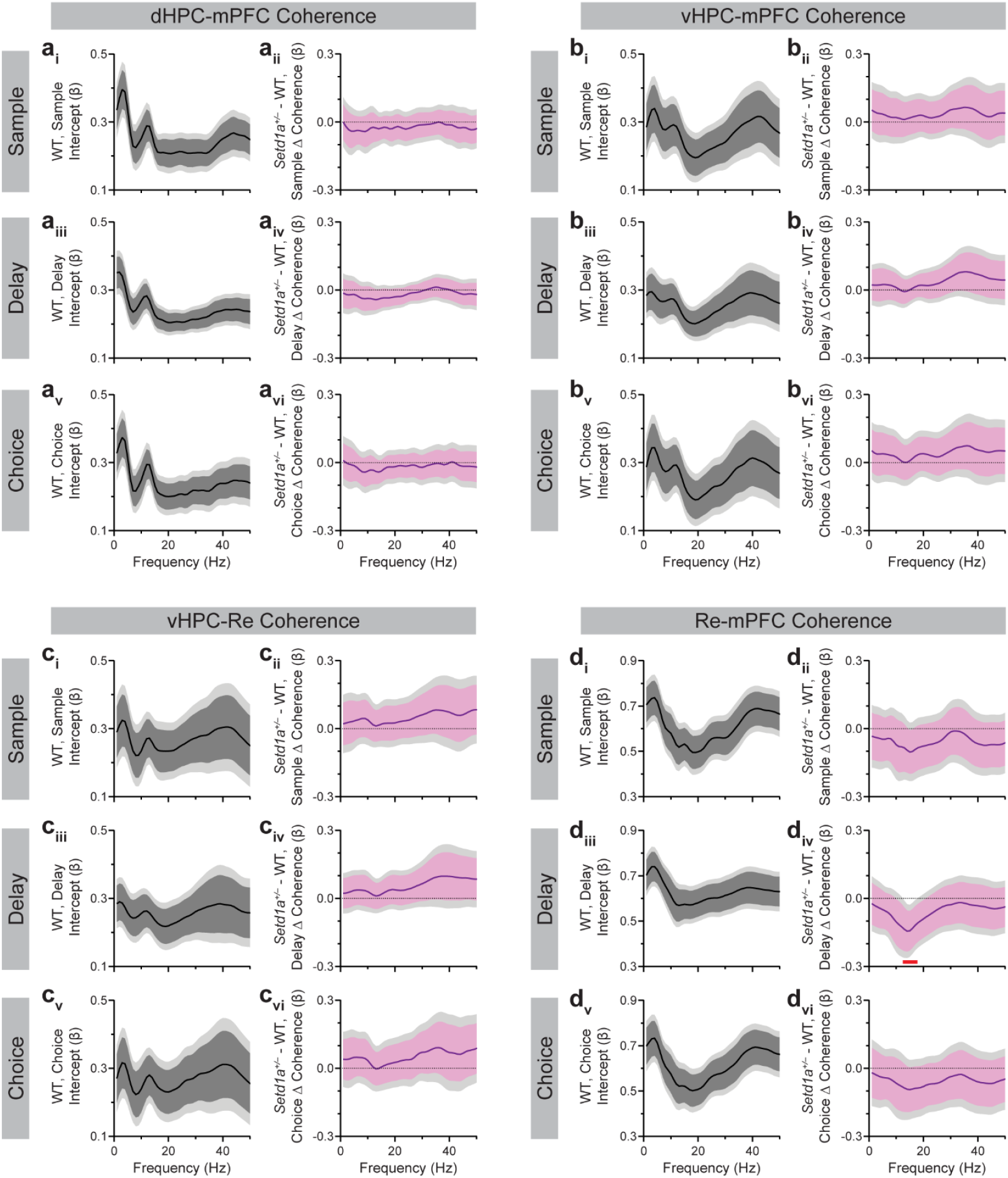
*Setd1a*^*+/–*^ mice show reduced Re-mPFC beta coherence during delay epoch of SWM task. (**a-d**) Functional coefficient estimates of the Genotype covariate from FLMM of epoch-level coherence for dHPC-mPFC (**a**), vHPC-mPFC (**b**), vHPC-Re (**c**), and Re-mPFC (**d**) brain region pairs for the sample (**i, ii**), delay (**iii, iv**), and choice (**v, vi**) epochs. Functional intercept estimates (**i, iii, v**) and Genotype coefficient estimates (**ii, iv, vi**) are presented for each region pair. Genotype coefficient estimates show the difference in average coherence at each frequency between *Setd1a*^*+/–*^ and WT mice. Pointwise and joint 95% confidence intervals (CI) are shown. Significant reductions (joint CIs do not contain zero) at specific frequencies are denoted by a red bar.

## Notes

### Competing Interest Statement

The authors have declared no competing interest.

